# Monosodium Iodoacetate delays regeneration and inhibits hypertrophy in skeletal muscle cells *in vitro*

**DOI:** 10.1101/577866

**Authors:** Rowan P. Rimington, Darren J. Player, Neil R.W. Martin, Mark P. Lewis

## Abstract

**Objective:** Osteoarthritis (OA) is a musculoskeletal disease which contributes to severe morbidity. The monosodium iodoacetate (MIA) rodent model of OA is now well established, however the effect of MIA on surrounding tissues post injection has not been investigated and as such the impact on phenotypic development is unknown. The aim of this investigation was to examine the impact of MIA incubation on skeletal muscle cells *in vitro*, to provide an indication as to the potential influence of MIA administration of skeletal muscle *in vivo*.

**Methods:** C2C12 skeletal muscle myotubes were treated with either 4.8μM MIA or 10μM Dexamethasone (DEX, positive atrophic control) up to 72hrs post differentiation and sampled for morphological and mRNA analyses.

**Results:** Significant morphological effects (fusion index, number of myotubes and myotube width, p<0.05) were evident, demonstrating a hypertrophic phenotype in control (CON) compared to a hyperplasic phenotype in MIA and DEX. Increases in MAFbx mRNA were also evident between conditions, with post-hoc analysis demonstrating significance between CON and DEX (p<0.001), but not between CON and MIA (p>0.05).

**Conclusions:** These data indicate a significant impact of both DEX and MIA on regeneration and hypertrophy *in vitro* and suggest differential activating mechanisms. Future investigations should determine whether skeletal muscle regeneration and hypertrophy is affected in the *in vivo* rodent model and the potential impact this has on the OA phenotypic outcome.

## 1. Introduction

Osteoarthritis (OA) is a debilitating musculoskeletal disease and presents a significant clinical problem in aging populations. OA manifests with pathological features including articular cartilage degradation, synovial inflammation, osteophyte formation and bone marrow lesions [1]. As the aetiology of OA is yet to be fully elucidated, *in vivo* pre-clinical animal models, which display comparable features to human OA, are particularly important in the basic and interventional research to understand disease onset, progression and potential therapeutic strategies [2].

One such model is the rodent mono-sodium iodoacetate (MIA) model of OA [3], which is now extensively published in the literature. Upon injection into the intra-articular space of rodent knee joints, MIA causes rapid articular cartilage damage and degeneration, leading to the development of an arthritic phenotype. The histopathology including cartilage lesion developments and proteogylcan matrix loss, along with functional joint impairment have all been shown to present in similar capacities to human OA [4–6]. Despite being an established pre-clinical *in vivo* model of OA, there is a paucity of research investigating the physiological implications of MIA administration on surrounding musculoskeletal tissues. Although MIA is injected into the avascular intra-articular space; as a small cell permeable molecule MIA has the potential to diffuse across the synovium initiating its direct interaction with other musculoskeletal tissues.

Skeletal muscle, of critical importance in OA pathology, plays an essential role in the stability of a joint when loading [7], and has been shown to directly regulate cartilage gene expression of collagen II and collagen IX proteins *in vitro* [8]. Furthermore, it is now clear that skeletal muscle size and strength contribute to a reduced risk of developing knee OA [9] and that skeletal muscle weakness and atrophy contribute to OA development and phenotype [10,11], particularly following previous musculoskeletal injury and arthroplasty [12]. Additionally, data suggest that quadriceps muscle weakness contributes to changes in key anabolic and catabolic matrix-regulating mRNA’s within key structural tissues of the joint (patellar tendon, medial and lateral collateral ligament, and medial and lateral meniscus [13]), indicating significant tissue interaction within the musculoskeletal system.

It is of paramount importance that debilitating changes in skeletal muscle phenotypes observed in pre-clinical animal models are entirely representative of the unloading/disuse/dysfunction associated with *in vivo* OA pathology, if such models are to have utility in understanding OA as an integrated musculoskeletal disease. As such, the degree to which MIA affects the surrounding tissues upon injection warrants investigation, to provide evidence as to whether the observed phenotype is developed through other means beyond primary cartilage degeneration. To this end, the aim of the current investigation was to develop a proof-of-concept model to investigate the effect of MIA on skeletal muscle cells *in vitro*.

## 2. Materials and methods

### 2.1. Cell culture

C2C12 murine myoblasts (ECACC, Sigma, UK) were used for all experimentation, and cultured in T80 flasks (Nunc™, Fisher Scientific, UK) maintained in a humidified 5% CO_2_ incubator (HERAcell 240i, Thermo Fisher, UK). Cells were maintained in growth media (GM) consisting of high glucose DMEM (Fisher Scientific) supplemented with 20% FBS (Dutscher Scientific, UK), 100U/ml Penicillin and 100µg/ml Streptomycin (Fisher Scientific) until 80% confluent, at which point cells were enzymatically detached from the flasks for either further sub-culture or experimentation. All experiments were conducted using cells below passage 13.

### 2.2. Experimental treatments

Myoblasts were seeded into 6 well plates at a density of 10 × 10^4^ cells/cm^2^ and cultured in GM until cells reached confluency. At this point cells were transferred to differentiation media (DM) consisting of high glucose DMEM (Fisher Scientific) supplemented with 2% Horse Serum (Dutscher Scientific), 100U/ml Penicillin and 100µg/ml Streptomycin (Fisher Scientific) for 72hrs to induce myotube formation. After 72hrs in DM and following 4hrs serum starvation, myotubes were treated with 10μM Dexamethasone (DEX, atrophic control) or 4.8μM MIA or control (CON) media over a time-course of 0, 6, 24 and 48hrs. A preliminary experiment was conducted to investigate a dose of MIA that would elicit an atrophic and myogenic response, without causing overt cell death (data not shown). Cells were fixed for morphological analyses and harvested for protein and PCR assays at the experimental time-points defined. Time point 0hrs was defined as 30min post transfer to experimental conditions.

### 2.3. Protein and mRNA analyses

For assessment of total cellular protein content, protein was extracted using 0.2M Sodium Hydroxide and protein concentrations were determined using the Pierce™ 660 protein assay (Fisher Scientific), according to manufactures instructions. To investigate the expression of putative atrophic genes, RNA was extracted using the TRIzol method, according to the manufacturer’s instructions (Sigma). mRNA expression of MuRF-1 and MAFbx were analysed as previously described with slight modifications [14]. 10μL volume PCR assays were conducted using Quantifast One Step RT-PCR kit (Qiagen, UK) on a ViiA7™ Real-Time PCR System (Applied bio-systems, Life Technologies). Data was analysed using the 2^(−ΔΔCT)^ method using POLR2B as a housekeeping gene with all data calibrated to a single CON 0hrs sample.

### 2.4. Fluorescence staining and microscopy

At the designated time-point cells were fixed and stained using F-actin molecular probe Rhodamine Phalloidin (Life Technologies, Molecular Probes) and counterstained with DAPI to visualise nuclei. Images were captured using a Leica DM2500 Fluorescent microscope at 20x magnification and analyses were conducted using Image J software.

### 2.5. Statistical analyses

All data are presented as mean ± SD. Differences between experimental conditions throughout the time-course investigated were analysed using a two-way ANOVA, using a Bonferroni Post-hoc test to analyse specific differences between isolated variables. All statistical analyses were conducted using SPSS version 22.0.

## 3. Results

### 3.1. MIA and DEX treatment affect myotube morphology

Various myogenic and morphological parameters were analysed to investigate the effects of MIA and DEX. Investigations of fusion index revealed a significant statistical main effect for time (p<0.001, figure 1b), suggesting the cells continue to differentiate throughout the experimental time-course. Despite no main effect evident for condition (p=0.439), a significant interaction effect between condition and time was found (p=0.018, figure 1b). These data suggest there is differential fusion index between conditions throughout the time-course investigated, providing evidence that both MIA and DEX contribute to delayed but potentiated myoblast fusion compared to CON at 48hrs.

**Figure 1.**
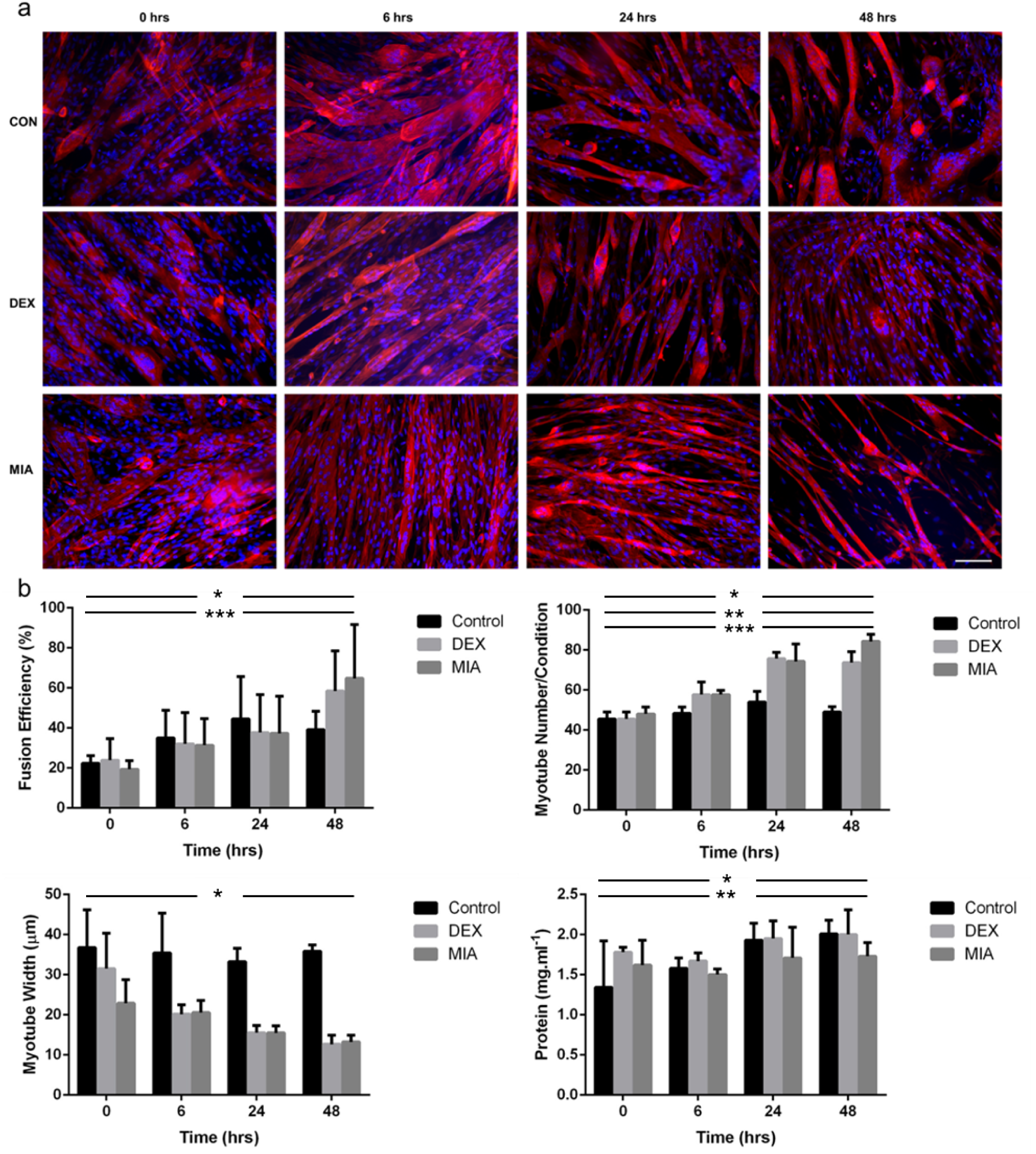
MIA induces significant morphological effects on skeletal muscle myotubes. (a) Cytochemical analyses demonstrate delayed regeneration and reduced hypertrophy in both DEX and MIA conditions compared to CON over a 48 hr experimental time-course. Red = Phalloidin, Blue = DAPI, Scale bar = 50 µm. (b) Quantitative analyses of fusion index revealed a significant statistical main effect for time (*p<0.001), no main effect for condition (p=0.439) and a significant interaction effect between condition and time (***p=0.018). Significant main effects for time (*p<0.001), condition (**p<0.001) and an interaction effect (***p<0.01) were evident for the number of myotubes. A significant main effect for condition (*p<0.001) and approaching significant effect for time (p=0.078) were found for myotube width. Significant differences in total cellular protein with main effects for condition (*p=0.034) and time (**p<0.001) despite no interaction effect (p=0.94) were evident.

Furthermore, significant main effects for time (p<0.001), condition (p<0.001) and an interaction effect (p<0.01, figure 1b) were evident for the number of myotubes. Post-hoc analyses demonstrated a greater number of myotubes present in both MIA and DEX conditions compared to CON (both p<0.001, figure 1b), with differences at 24hrs and 48hrs compared to 0hrs and 6hrs (all p<0.001, figure 1b). These data would suggest that both MIA and DEX contribute to a hyperplasic phenotype compared to CON, at the cost of reduced but more nucleated myotubes.

To further investigate the effects of MIA on skeletal muscle cells, myotube width was analysed as an indication of the extent of hypertrophy following fusion. A significant main effect for condition (p<0.001, figure 1b) and approaching significant effect for time (p=0.078, figure 1b), with post-hoc analyses providing evidence that CON myotubes were significantly more hypertrophic than those in either MIA or DEX conditions (both p<0.001, figure 1b). This was supported by significant total cellular protein content data, whereby main effects for condition (p=0.034, figure 1b) and time (p<0.001, figure 1b) despite no interaction effect (p=0.94) were evident. Taking all these morphological data together, it would appear that MIA contributes to a hyperplasic phenotype compared to a hypertrophic phenotype observed in CON, similar to that evident in the positive atrophic DEX condition.

### 3.2. MuRF1 and MAFbx expression

MuRF-1 and MAFbx are E3 ubiquitin ligases highly expressed in muscle wasting conditions and commonly used as markers of atrophy, due to their role in skeletal muscle protein breakdown. There was no effect of condition on the expression of MuRF1 mRNA (p = 0.517, figure 2a), despite an effect of time (p = 0.005, figure 2a), with no interaction effect. Post-hoc analyses revealed a significant decrease in MuRF1 mRNA at 48hrs compared to 6hrs (p = 0.003), with no differences between other time-points (p>0.05), suggesting early activation of this gene. A significant increase in MAFbx was found between conditions (p<0.001, figure 2b), despite no effect of time (p = 0.902) or any interaction effect (p=0.353, figure 2B). Post-hoc analyses demonstrated no differences between CON and MIA (p = 0.678) in MAFbx mRNA expression, despite significant increases in the expression of this gene in DEX compared to both CON and MIA conditions (both p<0.001 figure 2b). These data together suggest there is transcriptional activation of a component of the ubiquitin proteasome system in the DEX condition, which may in part be responsible for the reduced myotube width. In contrast, it appears that MIA treatment at the experimental time-points investigated fails to activate the ubiquitin proteasome system, and the reduced myotube width in this condition may be mediated through a different pathway.

**Figure 2.**
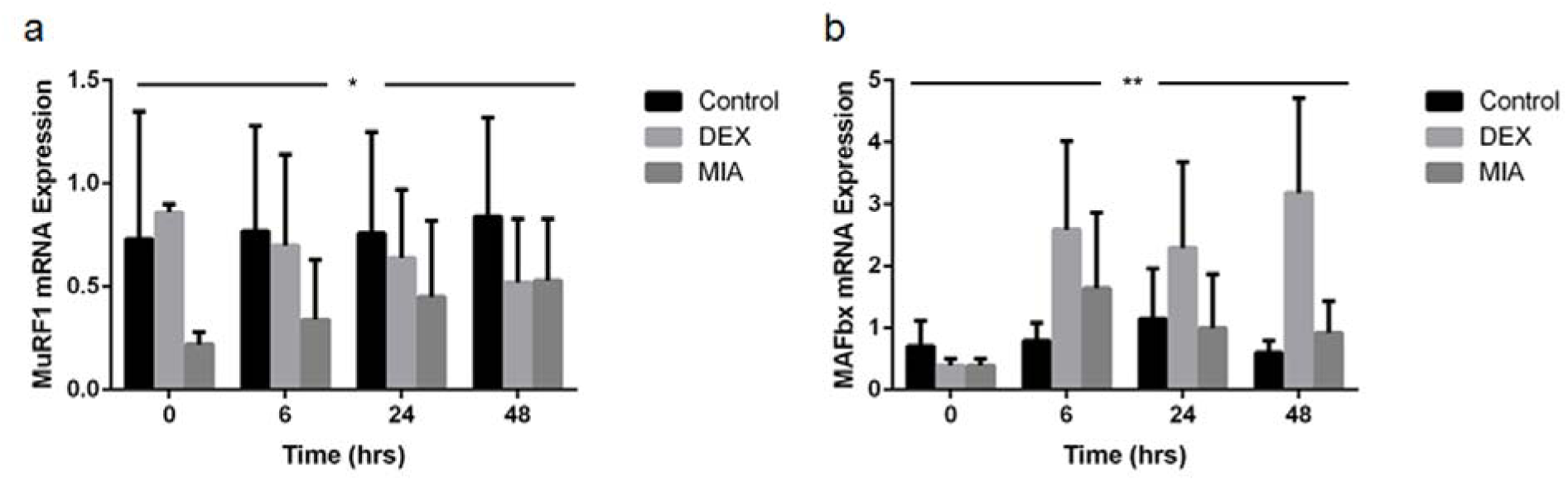
mRNA expression of members of the ubiquitin proteasome system. (a) No effect of condition on the expression of MuRF1 mRNA (p = 0.517), despite an effect of time (*p = 0.005), with no interaction effect. (b) A significant increase in MAFbx was found between conditions (**p<0.001, figure 2b), despite no effect of time (p = 0.902) or any interaction effect (p=0.353, figure 2b).

## 4. Discussion

Animal models serve as important tools in understanding the basic aetiology of OA and provide a necessary step in pre-clinical investigations of potential therapeutic strategies. A number of animal OA models have now been established including the MIA model. To date, the potential systemic influences of MIA in this model have not been fully investigated. Despite assuming that there are relatively low circulating concentrations of MIA, the deleterious effects to surrounding musculoskeletal tissues are yet to be investigated and hence the influence on the phenotypic outcome is currently unknown.

Here we sought to determine the effects of MIA on skeletal muscle cells and have used an *in vitro* model to demonstrate significant myogenic effects. Responses observed at early time-points were similar to that previously reported with DEX treatment; reduced differentiation and an apparent rapid atrophic phenotype [15,16]. Myotube width analyses confirmed a hypertrophic phenotype in CON compared to MIA and DEX, however, only with DEX treatment was there an observed increase in MAFbx mRNA, supporting previous findings for its atrophic role in this condition [17].

It is important to consider however the delayed differentiation and hyperplasic phenotype in both MIA and DEX conditions at the later time-points, a phenotype suggested to be regulated by myogenic regulatory factors (MRF’s) myoD and myogenin [18]. This hyperplasic response, along with the apparent increases in fusion efficiency at 48hrs in DEX and MIA compared to CON, would suggest that both treatment conditions have the ability to stimulate and potentially augment myogenesis, albeit in a delayed capacity. An increased differentiation and hypertrophic capacity of C2C12’s treated with DEX when co-incubated with IGF-I, has previously been reported [19]. This is thought to occur through the IGF-I-meditated translation of accumulated muscle-specific mRNA transcripts induced by DEX [19] and may in part explain the delayed, yet augmented hyperplasic phenotype observed in the DEX condition presented here.

Despite similar morphological findings between MIA and DEX conditions, the primary activated mechanisms which contribute to this shared phenotype are potentially distinct. DEX-mediated skeletal muscle atrophy is thought to occur through the initiation of mechanisms regulating protein breakdown. Specifically, increases in MuRF1 and MAFbx mRNA’s following DEX has been attributed to the activation of FOXO1 and FOXO3A, as a result of reduced Akt activity [17]. The mechanism regulating the hyperplasic and reduced myotube width phenotype observed in the MIA condition is currently unknown, however given the glycolytic inhibitory capacity of MIA, it could be speculated that this response is a consequence of disrupted cellular energy status. Increases in AMPK activity as a consequence of an augmented AMP-to-ATP ratio, has been shown to inhibit mTOR, through the phosphorylation of TSC2 [20], inhibiting the key mechanism for protein synthesis and hence reducing the ability for the differentiated myotubes to hypertrophy.

The interaction between cartilage and skeletal muscle tissue in OA pathology is critical to the development of reproducible pre-clinical models. *In vivo*, primary cartilage degradation contributes to significant biomechanical changes and is thought to affect skeletal muscle through pain and limb unloading mechanisms contributing to fibre atrophy and subsequent weakness. Reduced myogenic regeneration and inhibition of skeletal muscle hypertrophy observed in this work indicates that should MIA become systemic, deleterious effects on the surrounding tissue may well be apparent and significant.

## 5. Conclusions

We have demonstrated significant myogenic effects of MIA in an isolated *in vitro* model, similar to that seen in DEX. Future work should seek to determine if MIA administration affects skeletal muscle regeneration and hypertrophy *in vivo* and in particular whether such potential negative outcomes, contribute to the development of the observed OA phenotype. This will need to be investigated using an integrated approach of *in vivo* testing of muscle function and a raft of *ex vivo* analyses to understand any impact at the functional and molecular levels respectively.

